# *Rhopalocnemis phalloides* has one of the most reduced and mutated plastid genomes known

**DOI:** 10.1101/448787

**Authors:** Mikhail I. Schelkunov, Maxim S. Nuraliev, Maria D. Logacheva

**Affiliations:** Skolkovo Institute of Science and Technology, Moscow, Russia; Institute for Information Transmission Problems, Moscow, Russia; Faculty of Biology, Lomonosov Moscow State University, Moscow, Russia; Joint Russian–Vietnamese Tropical Scientific and Technological Center, Cau Giay, Hanoi, Vietnam; A.N. Belozersky Research Institute of Physico-Chemical Biology, Lomonosov Moscow State University, Moscow, Russia

## Abstract

Although most plant species are photosynthetic, several hundred species have lost the ability to photosynthesize and instead obtain nutrients via various types of heterotrophic feeding. Their genomes, especially plastid genomes, markedly differ from the genomes of photosynthetic plants. In this work, we describe the sequenced plastid genome of the heterotrophic plant *Rhopalocnemis phalloides*, which belongs to the family Balanophoraceae and feeds by parasitizing on other plants. The genome is highly reduced (18 622 base pairs versus approximately 150 kilobase pairs in autotrophic plants) and possesses an outstanding AT content, 86.8%, the highest of all sequenced plant plastid genomes. The gene content of this genome is quite typical of heterotrophic plants, with all of the genes related to photosynthesis having been lost. The remaining genes are notably distorted by a high mutation rate and the aforementioned AT content. The high AT content has led to sequence convergence between some of the remaining genes and their homologues from AT-rich plastid genomes of protists. Overall, the plastid genome of *R. phalloides* is one of the most unusual plastid genomes known.

## Introduction

Though plants are generally considered photosynthetic organisms, there are several hundred plant species that have lost the ability to photosynthesize during the course of evolution (Westwood et al., 2010; Merckx, 2013). They feed by parasitizing either on fungi or on other plants. In addition to the completely heterotrophic plants, there are also plants that combine the ability to photosynthesize with the heterotrophic lifestyle. They are termed partial heterotrophs (or hemi-heterotrophs, or mixotrophs) in contrast to the former, which are termed complete heterotrophs (or holo-heterotrophs).

The completely heterotrophic plants show a high degree of similarity, though there were several dozen cases of independent transition to complete heterotrophy. For example, these plants all either lack leaves or have very reduced leaves. These plants are non-green because of the absence (or at least highly reduced amounts (Cummings and Welschmeyer, 1998)) of chlorophyll. Additionally, a common feature of many completely heterotrophic angiosperms is that they spend most of their lifetimes underground, since without the need to photosynthesize their only reason to appear aboveground is for flowering and seed dispersal.

Genomic studies of heterotrophic plants are mostly focused on plastid genomes, since 1) most of the plastid genes are related to photosynthesis, and thus changes in the plastid genomes are expected to be more prominent compared to mitochondrial and nuclear genomes, and 2) plastid genomes are smaller than nuclear and mitochondrial ones and usually have higher copy numbers and are thus easier to sequence (Daniell et al., 2016; Gualberto and Newton, 2017; Sakamoto and Takami, 2018). The main feature of the plastid genomes of complete heterotrophs is the loss of genes responsible for photosynthesis and respective shortening of the genomes, from approximately 150 kbp (typical of autotrophic plants) to, in the most extreme known case, 12 kbp (Bellot and Renner, 2015; Wicke and Naumann, 2018). The remaining genes are the ones with functions not related to photosynthesis. Usually they are *accD* (a gene whose product participates in fatty acid synthesis—one of the plastid functions besides photosynthesis), *clpP* (encodes a component of a complex responsible for degradation of waste proteins in plastids), *ycf1* (encodes a component of the translocon—the complex which imports proteins from cytoplasm into plastids), *ycf2* (a conserved gene present in almost all plants, but with unknown function) and various genes required for translation of the aforementioned ones, namely genes which code for protein and RNA components of the plastid ribosome and for tRNAs. One of the tRNA-coding genes, *trnE-UUC*, has also an additional function, with its product participating in haem synthesis (Kumar et al., 1996).

In addition to the expected shortening of the genome, there are some peculiar and still unexplained features in the plastid genomes of heterotrophic plants, namely their increased mutation accumulation rate (Bromham et al., 2013; Wicke and Naumann, 2018) and increased AT content (Wicke and Naumann, 2018). In the most extreme cases, plastid genomes of heterotrophic plants may accumulate mutations approximately 100 times faster than their closest autotrophic relatives (Bellot and Renner, 2015). The most obvious explanation, the relaxation of selection, is refuted by the fact that dN/dS (a common measure of selective pressure) is usually not increased in the plastid genes of heterotrophic plants, except for photosynthesis-related genes during their pseudogenization (Logacheva et al., 2011; Barrett et al., 2014; Schelkunov et al., 2015; Lam et al., 2015; Wicke et al., 2016; Naumann et al., 2016). AT content is increased from circa 65% in autotrophic species (Smith, 2012) to 77% in the most prominent case among heterotrophic species (Naumann et al., 2016), also for an unknown reason.

Genes not related to photosynthesis, such as *accD* and *infA*, are sometimes transferred to the nuclear genome (Millen et al., 2001; Rousseau-Gueutin et al., 2013; Liu et al., 2016). Therefore, when all genes with functions not related to translation are transferred to the nuclear genome, there will be no reasons to keep the translation apparatus in plastids, and the genes responsible for translation will also be lost. Thus, the plastid genome is potentially able to disappear entirely. Indeed, two cases of the complete plastid genome loss are known: one in algae of the genus *Polytomella* (Smith and Lee, 2014) and one in the parasitic plant *Rafflesia lagascae* (Molina et al., 2014).

The initial aim of the present study was to prove that a completely heterotrophic plant, *Rhopalocnemis phalloides*, had also lost its plastid genome completely. *Rhopalocnemis phalloides* is a parasitic plant from the family Balanophoraceae (order Santalales) which occurs in Asia and feeds by obtaining nutrients from roots of various plants. Initially we sequenced circa 10 million pairs of reads on the HiSeq 2000 platform and observed no contigs with similarity to typical plastid genes, while there were obvious mitochondrial contigs. Based on our experience in studying plastid genomes of heterotrophic plants, mitochondrial contigs usually have lower sequencing coverage than plastid ones, and thus the plastid genome is always easier to assemble. This led us to suppose that the plastid genome in *R. phalloides* may have been lost completely. To verify this, we sequenced approximately 200 million pairs of additional reads. What we found is that the plastid genome *is* in fact present, but its tremendous AT content (86.8%) hampered PCR, which is one of the usual steps in library preparation of Illumina, and thus the sequencing coverage of the genome was much lower than one might have expected. This article is dedicated to the analysis of this plastid genome.

## Materials and Methods

### Sample collection and sequencing

The specimen of *R. phalloides* was collected during an expedition of the Russian-Vietnamese Tropical Centre in Kon Tum province, Vietnam, in May 2015 (voucher information: Southern Vietnam, Kon Tum prov., Kon Plong distr., Thach Nham protected forest, 17 km N of Mang Den town, in open forest, N 14° 45′ 15′′ E 108° 17′ 40′′, elev. 1400 m, Nuraliev M.S., Kuznetsov A.N., Kuznetsova S.P., No. 1387, 18.04.2015). The studied material was preserved in silica gel and in RNAlater. The voucher is deposited at the Moscow University Herbarium (MW) (Seregin, 2018) with the barcode MW0755444.

DNA was extracted from an inflorescence using a CTAB-based method (Doyle, 1987), and the DNA library was prepared using the NEBNext DNA Ultra II kit (New England Biolabs). Sequencing was performed with a NextSeq 500 sequencing machine (Illumina) in the paired end mode, producing 387 351 294 reads (193 675 647 read pairs), each 150 bp long.

RNA was extracted from an inflorescence using the RNeasy Mini kit (Qiagen). Plastid transcripts are usually not polyadenylated, so the method of poly(A) RNA selection was not applicable in our study. Instead, we used a protocol based on depletion of ribosomal RNA with the Plant Leaf Ribo Zero kit (Illumina). The RNA-seq library was prepared using the NEBNext RNA Ultra II kit (New England Biolabs) and sequenced on a HiSeq 2500 sequencing machine (Illumina) with TruSeq reagents v.4 in the paired end mode, producing 54 794 466 reads (27 397 233 read pairs), 125 bp each.

### Genome assembly and annotation

Both DNA-seq and RNA-seq reads were trimmed by Trimmomatic 0.32 (Bolger et al., 2014) in the palindromic mode, removing bases with quality less than 3 from the 3′ ends of reads, and fragments starting from 4-base-long windows with average quality less than 15 (SLIDINGWINDOW:4:15). Reads which, after trimming, had average quality less than 20 or length shorter than 30 bases were removed.

The assembly was performed from DNA-seq reads by two tools. First, it was made by CLC Assembly Cell 4.2 (https://www.qiagenbioinformatics.com/products/clc-assembly-cell/) with the default parameters. Second, it was made by Spades 3.9.0 (Bankevich et al., 2012). As Spades performs slowly when running on large number of reads, prior to starting its assembly we removed from reads k-mers with coverage less than 50× by Kmernator 1.2.0 (https://github.com/JGIBioinformatics/Kmernator). This allowed us to eliminate most reads belonging to the nuclear genome (and, potentially, some reads belonging to low-covered plastid regions), thus highly reducing the number of reads. The Spades assembly was run on this reduced read set, with the “--only-assembler” and “--careful” options. To determine the read coverage of contigs in these two assemblies, we aligned to them reads by CLC Assembly Cell 4.2, requiring at least 80% of the length of each read to align with a sequence similarity of at least 98%.

To find contigs potentially belonging to plastid and mitochondrial genomes, we aligned by BLASTN and TBLASTN from BLAST 2.3.0+ suit (Camacho et al., 2009) proteins and non-coding RNA (ncRNA) genes from reference species. As the references, we used sequences from the plastid genomes of *Viscum album* (NCBI accession NC_028012), *Osyris alba* (NCBI accession NC_027960), *Arabidopsis thaliana* (NCBI accession NC_000932), *Nicotiana tabacum* (NCBI accession NC_018041) and mitochondrial genomes of *Viscum album* (NCBI accession NC_029039), *Citrullus lanatus* (NCBI accession GQ856147), *Mimulus guttatus* (NCBI accession NC_018041). *Viscum album* and *O. alba* were used because they, like *R. phalloides*, belong to Santalales. Other species were chosen because they belong to other orders of eudicots. Alignment was performed with the maximum e-value of 10^-3^ and low complexity filter switched off. The word size was 7 for BLASTN and 3 for TBLASTN. Here and later, the local BLAST was used with the parameter “max_target_seqs” set to 10^9^ to avoid the problem discussed by Shah et al. (2018).

Five contigs containing plastid genes were found in the CLC assembly and 3 contigs in the Spades assembly. After aligning contigs of these two assemblies to each other (BLASTN, maximum e-value 10^-3^, word size 7, low complexity filter switched off), it appeared that places where CLC contigs were broken by gaps corresponded to continuous places in the Spades contigs and, vice versa, gaps in the Spades contigs corresponded to continuous places in the CLC contigs. This allowed us, by joining the contigs of these two assemblies, to create a circular sequence corresponding to the plastid genome. To check the assembly, we mapped reads (in CLC Assembly Cell 4.2, requiring at least 80% of the length of each read to align with a sequence similarity of at least 98%) to the resultant sequence and verified (by eye, in CLC Genomics Workbench 7.5.1, https://www.qiagenbioinformatics.com/products/clc-genomics-workbench/) that there were no places uncovered by reads and no places where the insert size abruptly decreased or increased. Such places of abrupt increase or decrease of the insert size may indicate regions with assembly errors, consisting of, respectively, sequence insertions or deletions. As read mapping is complicated on the edges of a sequence, we also performed such analysis on a reoriented version of the plastid genome, where the sequence was broken in the middle and the ends were joined. These analyses indicated that the assembly contained no errors.

To find genes in the plastid genome, we used a complex strategy, since highly mutated genes may be hard to notice. We used the following methods:

1. The alignment by BLASTN and TBLASTN of reference protein-coding and ncRNA-coding genes, described above.
2. Open reading frames were scanned by InterProScan 5.11 (Jones et al., 2014) using the InterPro 51.0 (Finn et al., 2017) database with the default parameters. “Open reading frames” here were any sequences at least 20 codons long uninterrupted by stop codons. Not requiring an ORF to begin from a start-codons allowed for the detection of exons in multi-exonic genes.
3. The genome was scanned by Infernal 1.1.2 (Nawrocki and Eddy, 2013) with RNA models from Rfam 12.2 database (Nawrocki et al., 2015) to predict ncRNA-coding genes. The maximum allowed e-value was set to 10^-3^.
4. To predict rRNA-coding genes, RNAmmer 1.2 server (Lagesen et al., 2007) was used in bacterial mode and eukaryotic mode.
5. The genome was scanned by tRNAscan-SE 1.23 (Lowe and Eddy, 1997) with the default parameters, in the organellar (models trained on plastid and mitochondrial tRNAs) and also in the general (models trained on tRNAs from all three genomes) mode, to predict tRNA-coding genes.
6. The genome was annotated by DOGMA (Wyman et al., 2004) and Verdant (McKain et al., 2017).
7. When determining which ATG codon was a true start codon, we compared the sequence of a gene with sequences of its homologs from the aforementioned reference species.
8. To determine exon borders, RNA-seq reads with minimal length of 100 bp (to minimize false mappings) were mapped to the genome by CLC Assembly Cell 4.2, requiring at least 50% of each read’s length to map with a sequence similarity of at least 90%. Exon borders were found by eye in CLC Genomics Workbench 7.5.1 as regions of genes where many reads were mapped partially. The exon borders of the reference species were used for comparison.
9. To check for RNA editing that could create new start or stop codons, we mapped RNAseq reads with a minimal length of 100 bp by CLC Assembly Cell 4.2, requiring at least 80% of each read’s length to map with a sequence similarity of at least 90%. Mismatches between the reads and the genome were inspected by eye in CLC Genomics Workbench 7.5.1.
10. After annotating the genes, we additionally verified that there were no remaining regions with high sequence complexity, relatively low AT content or high coverage by RNA-seq reads where no genes were predicted. Regions with high sequence complexity were predicted in the genome by CLC Genomics Workbench 7.5.1 using K2 algorithm (Wootton and Federhen, 1993) with a window size of 101 bp. The AT-content plot was created by a custom script with 200-bp-long windows. RNA-seq reads with a minimal length of 100 bp were mapped by CLC Assembly Cell 4.2, requiring at least 80% of each read’s length to map with a sequence similarity of at least 90%.

After completing gene prediction, the plastid genome was reoriented to start from the first position of *rps14*, as this is the first gene in the canonical representation of the plastid genome of *A. thaliana* which is also present in the plastid genome of *R. phalloides*.

### Estimation of contamination amount

To estimate the amount of contamination, 1000 random DNA-seq read pairs, taken after the trimming, were aligned by BLAST to NCBI databases. Taxonomies of their best matches were used as proxies for the reads’ source taxonomies. To increase the sensitivity of the search, the analysis was performed as follows:

1. All reads were aligned to NCBI NT (the database current as of September 18, 2017) by BLASTN from BLAST 2.3.0+ suite with the maximum allowed e-value of 10^-3^ and the word size of 7 bp. To decrease the number of false-positive matches, hard masking of low-complexity regions (“-soft_masking false” option) was used.
2. All reads were aligned to NCBI NR (the database current as of September 18, 2017) by BLASTX from BLAST 2.3.0+ suite with the maximum allowed e-value of 10^-3^ and the word size of 3 bp. Hard masking in BLASTX is enabled by default.
3. If at least one of two reads in a pair had matches to NT, the taxonomy of the match with the lowest e-value was considered the taxonomy of the read pair. If the read pair had no matches in NT, the taxonomy of the match to NR with the lowest e-value was considered the taxonomy of the read. Thus, the alignment to NT had higher priority than the alignment to NR. This was done to take into account synonymous positions of genes, where possible, and thus increase the precision of the taxonomic assignment of read pairs.

### Other analyses

To determine the phylogenetic placement of *R. phalloides* within Balanophoraceae, we utilized the alignment of genes from 186 species (180 species of Santalales plus 6 outgroup species) created by Su et al. (2015). *Rhopalocnemis phalloides* was not studied in that article. Seven genes were used for the phylogenetic analysis in that work: plastid *accD*, *matK*, *rbcL*; nuclear 18S rDNA, 26S (also known as 25S) rDNA and *RPB2*; and mitochondrial *matR*. As *matK* and *rbcL* are absent from the plastid genome of *R. phalloides*, we were unable to use them. *accD* of *R. phalloides* contains many mutations and thus can be aligned improperly, so we did not use it either. Mitochondrial *matR* is disrupted in *R. phalloides* by several frameshifting indels. Due to the large size of the nuclear genome of *R. phalloides* (see the paragraph “Other genomes of *R. phalloides*”), *RPB2* had low coverage, and its sequence could not be obtained from the available DNA-seq reads. The sequences of 18S rDNA and 26S rDNA were easier to determine, as they had many copies in the nuclear genome and thus their coverage was higher. To find their sequences among the contigs, we aligned 18S rDNA and 26S rDNA of *A. thaliana* by BLASTN with the default parameters to the contigs of the Spades assembly. The sequences of 18S rDNA and 26S rDNA were added to the alignment of Su et al. (2015) using MAFFT 7.402 (Katoh and Standley, 2013) with options --addfragments and --maxiterate 1000. The phylogenetic tree was built with RAxML 8.2.4 (Stamatakis, 2014), utilizing 20 starting stepwise-addition parsimony trees, employing GTR+Gamma model, with the same 6 outgroup species as in the work of Su et at. (2015) (*Antirrhinum majus*, *A. thaliana, Camellia japonica*, *Cornus florida*, *Myrtus communis* and *Spinacia oleracea*). The required number of bootstrap pseudoreplicates was determined by RAxML automatically with the extended majority-rule consensus tree criterion (“autoMRE”). The tree was visualized with FigTree 1.4.3 (http://tree.bio.ed.ac.uk/software/figtree/).

To compare the substitution rate in the plastid genome of *R. phalloides* with substitution rates in other species of Santalales, we used common protein-coding genes of *R. phalloides*, *A. thaliana* (used as the outgroup) and the only 4 species of Santalales with published plastid genomes as of 2017: *V. album*, *O. alba*, *Champereia manillana* and *Schoepfia jasminodora*. Their protein-coding gene alignment was created by TranslatorX 1.1 (Abascal et al., 2010) based on an alignment of corresponding amino acid sequences performed by Muscle 3.8.31 (Edgar, 2004) with the default parameters. *ycf1*, *ycf2* and *accD* of *R. phalloides* differed from homologous genes of other species so much that a reliable alignment was not possible. Alignments of other genes were then concatenated into a single alignment and provided to Gblocks server (Castresana, 2000), which removed poorly aligned regions from the alignment. Gblocks was run in the codon mode, with the default parameters. Substitution rates and selective pressure were evaluated by codeml from PAML 4.7 (Yang, 2007) with the F3×4 codon model, starting dN/dS value of 0.5 and starting transition/transversion rate of 2. The phylogenetic tree provided to PAML was a subtree of the large phylogenetic tree of Santalales, produced as described above. Additionally, the analysis of substitution rates and selective pressure was performed by BppSuite 2.3.2 (Guéguen et al., 2013). To our knowledge, this is the only tool that is capable of phylogenetic analyses of protein-coding sequences which takes into account different codon frequencies in different sequences (Guéguen and Duret, 2017), while PAML uses a single averaged codon frequency for all sequences. This is important, as the codon frequencies in *R. phalloides* highly differ from the codon frequencies in the mixotrophic Santalales of comparison (see the Supplementary Table 2). The program bppml from BppSuite was run using a nonhomogeneous (“one_per_branch”) model, the substitution model was YN98, the codon model F3×4, starting dN/dS values of 0.5 and starting transition/transversion rates of 2. Starting branch lengths were 0.1 substitutions per codon. The parameter estimation was performed by the full-derivatives method with optimization by the Newton-Raphson method (“optimization=FullD(derivatives=Newton)”), using parameters transformation (“optimization.reparametrization=yes”).

To check for similarity between the genes and the proteins of the *R. phalloides* plastid genome and sequences from other species, we performed BLASTN and BLASTP alignment against NCBI NT and NR databases, respectively, on the NCBI website (https://blast.ncbi.nlm.nih.gov/Blast.cgi) on March 4, 2018 with the default parameters.

To build the phylogenetic tree of *rrn16* (Figure 4A), sequences from *R. phalloides*, *O. alba*, *V. album*, *N. tabacum*, *A. thaliana* were supplemented with sequences from *Corynaea crassa* (NCBI accession U67744), *Balanophora japonica* (NCBI accession KC588390), *Nitzschia* sp. IriIs04 (NCBI accession AB899709), *Leucocytozoon caulleryi* (NCBI accession AP013071) and *Plasmodium cynomolgi* (NCBI accession AB471804). These particular sequences of *Nitzschia*, *Leucocytozoon* and *Plasmodium* were randomly chosen among the SAR sequences that produced matches to the *rrn16* of *R. phalloides* in the analysis described in the previous paragraph. The *rrn16* sequences were aligned by Muscle 3.8.31. Poorly aligned regions were removed by Gblocks server. The phylogenetic tree was built by RAxML 8.2.4 with the parameters described above. The phylogenetic tree of species in Figure 4A was taken from the TimeTree database (timetree.org). The subtree of Balanophoraceae in the tree was taken from the general tree of Santalales (Supplementary Figure 5), created as described above. The plastids of species from the SAR (“Stramenopiles, Alveolata, Rhizaria”) clade originated from a secondary endosymbiosis with a red alga, but this endosymbiosis occurred in SAR once in the root, and thus (taking into account that red algae is an outgroup to Embryophyta) the phylogenetic tree of plastids of the studied species coincide with the phylogenetic tree of nuclei. The trees were drawn by TreeGraph 2.14.0-771 beta (Stöver and Müller, 2010).

Codon usage and amino acid usage of the common protein-coding genes of *R. phalloides* and species of comparison were calculated by CodonW 1.4.2 (Peden, 1999). Frequencies of 21-bp-long k-mers were calculated for the trimmed DNA-seq reads by Jellyfish 2.1.2 (Marçais and Kingsford, 2011), not using the Bloom filter (to count the number of low-frequency k-mers precisely).

The list of plastid genomes with their lengths and AT contents was obtained from the NCBI database (https://www.ncbi.nlm.nih.gov/genome/browse/#!/organelles/). Information on whether a specific plant species is completely heterotrophic was obtained by literature analysis.

## Results and discussion

### The gene content of the *R. phalloides* plastid genome

The plastid genome of *R. phalloides* is a circular molecule 18 622 bp long. Its map is represented in Figure 1, in a linear form, for convenience. The protein-coding gene content is quite typical for highly reduced plastid genomes of completely heterotrophic plants (Wicke and Naumann, 2018). The plastid genome of *R*.*phalloides* possesses the genes *accD*, *clpP*, *ycf1*, *ycf2* (mentioned in the Introduction) and 9 genes encoding protein components of the ribosome. Additionally, it codes for *rrn16* and *rrn23*, RNA components of the plastid ribosome. Like several other highly reduced plastid genomes, it lacks *rrn4.5* and *rrn5*—genes coding for two other RNAs of the ribosome, which poses the interesting puzzle of how the ribosome works without these genes in the plastid genome. One possibility is that these genes were transferred either to the mitochondrial or to the nuclear genome and are now transcribed there and imported to the plastids from the cytoplasm. The other possibility is that the ribosome is capable of working without them, akin to how it can work without some ribosomal proteins (Tiller and Bock, 2014). We plan to clarify this question in an upcoming article dedicated to the analysis of the *R*. *phalloides* transcriptome.

**Figure 1.**
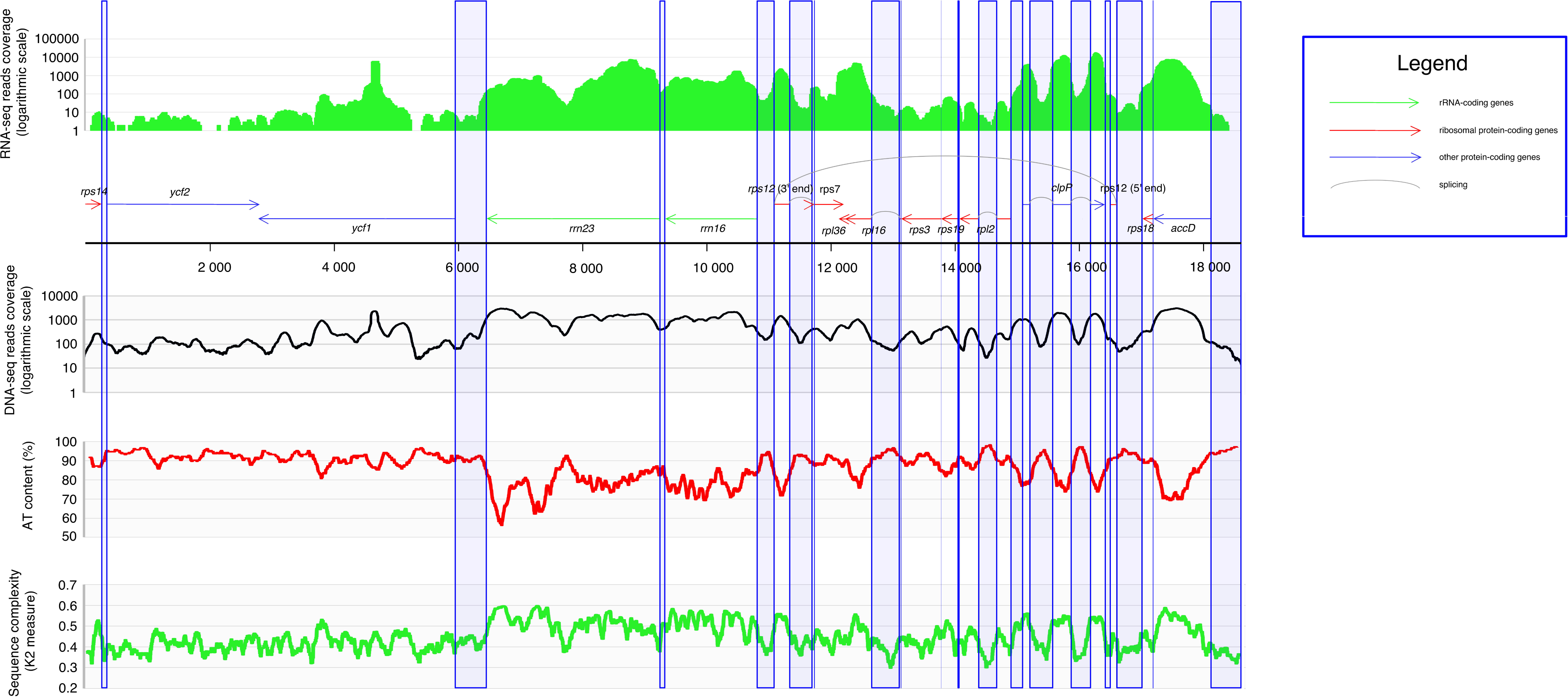
Map of the *Rhopalocnemis phalloides* plastid genome showing various features. The circular plastid genome is represented linearly for convenience. Blue columns denote non-coding regions.

The tRNA-coding gene content of the *R*. *phalloides* plastid genome also puzzles. The standard method to predict tRNA-coding genes is the program tRNAscan-SE. It has a dedicated “organellar” mode where tRNA models were trained on mitochondrial- and plastid-encoded tRNA sequences and structures. It also has a “general” mode whose models are based on nuclear-encoded tRNAs. In the organellar mode the tool predicts 64 tRNA-coding genes, which is much more than the approximately 30 tRNA-coding genes encoded in plastomes of typical autotrophic species (Wicke et al., 2011). In the general mode, the tool predicts zero tRNA-coding genes. Our experience in working with different plastid genomes suggests that results of predictions in these two modes usually coincide. Of the 64 predicted tRNA-coding genes, 61 have introns, and the mean AT content of the 64 genes is 94%. Thus, we suppose that most of them, if not all, are false-positive predictions. They may originate from the ease with which sequences of low complexity form secondary structures—these spuriously generated cloverleaf-like structures may have deceived the algorithms of tRNAscan-SE. Seventeen of the predicted tRNA-coding genes are for isoleucine tRNAs, and 11 are for lysine. This further attests to the false-positive nature of these genes, as false-positively predicted tRNA-coding genes in an AT-rich genome are expected to have AT-rich anticodons, and the anticodons of isoleucine and lysine tRNAs are two of the most AT-rich of all amino acid anticodons. Of the three tRNA-coding genes without introns, one has an AT content of 76%, another 92% and the third 96%. Because of the relatively low AT, the first seems to be a possible candidate for a true gene. Its AT content is not only the lowest among the three predictions which do not have introns but also among all 64 predicted tRNA-coding genes. This is a *trnL* gene with anticodon TAA (UAA). Nevertheless, we cannot tell confidently whether this gene is a false-positive prediction and thus did not use it for any analyses. The only predicted *trnE* gene has an AT content of 99% and is thus very likely to be a false prediction. Therefore, the plastid genome of *R. phalloides* has probably lost its *trnE*, like the plastid genomes of completely heterotrophic plants from the genus *Pilostyles* did (Bellot and Renner, 2015), although earlier *trnE* was deemed indispensable because of its function in haem synthesis (Barbrook et al., 2006; Howe and Smith, 1991). Potentially, *trnE* could have been transferred to the nuclear or the mitochondrial genome, transcribed there and imported into the plastids of *R. phalloides* from the cytoplasm.

### The plastid genome of *R. phalloides* has very high AT content

One of the most interesting features of the *R. phalloides* plastid genome is its AT content of 86.8%. It is the highest among all sequenced plant plastid genomes, the second being *Pilostyles hamiltonii* with 77.3% (Bellot and Renner, 2015). Among eukaryotes and prokaryotes, there are several known genomes with higher AT content, all belonging to mitochondria or apicoplasts, with the record held by the mitochondrial genome of a fungus, *Nakaseomyces bacillisporus* CBS 7720 (Bouchier et al., 2009), with the AT content of 89.1% (according to the NCBI site, information current as of June 14, 2018).

The increased AT content is a common feature of plastid genomes of completely heterotrophic plants (Figure 2, Supplementary Table 1). To our knowledge, it remains unexplained. It correlates with the degree of plastid genome reduction, with plastid genomes of plants such as *R. phalloides* or *Pilostyles hamiltonii* having simultaneously some of the largest AT contents and some of the smallest plastid genomes.

**Figure 2.**
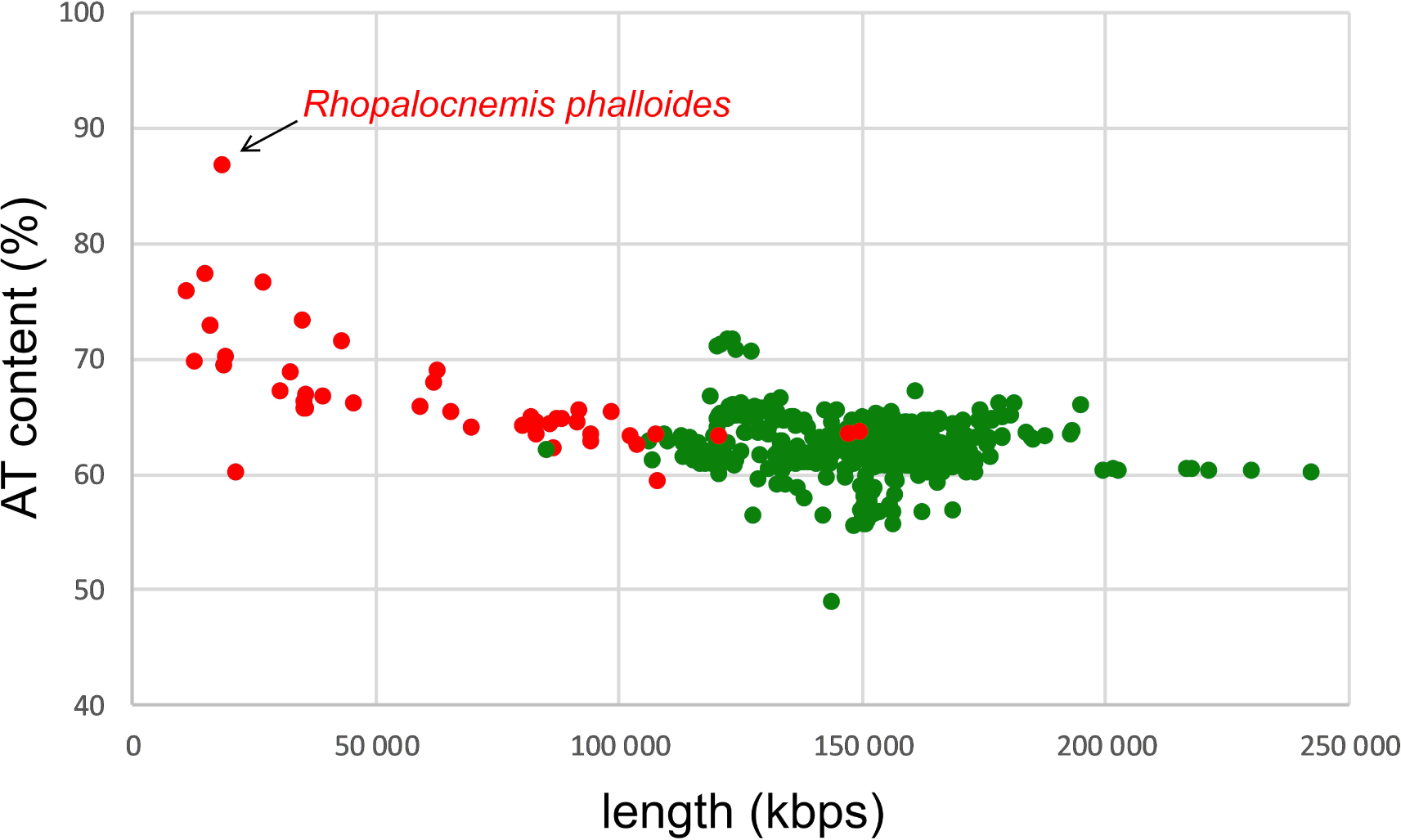
AT contents and lengths of the plastid genomes of Embryophyta. Red dots denote completely heterotrophic plants and green dots mixotrophic and completely autotrophic.

It was the high AT content which prevented us from detecting the plastid genome of *R. phalloides* from the initial assembly made of circa 10 million read pairs. High AT content hampers PCR (Benjamini and Speed, 2012), and as library preparation for Illumina sequencing machines usually involves PCR, coverage of AT-rich regions is decreased. The sequencing coverage in the *R. phalloides* plastid genome ranges from circa 3000× in the least AT-rich regions to 17 in the most AT-rich ones (Supplementary Figure 1). The AT content and the sequencing coverage correlate with a Spearman’s correlation coefficient of −0.93. Read insert size also depends on the AT content, with the least AT-rich regions covered by reads with an insert size of approximately 300 bp and the most AT-rich regions with an insert size of approximately 200 bp (Supplementary Figure 2); Spearman’s correlation coefficient is −0.69. We suppose that the coverage drop associated with high AT content may be the reason why the authors of a work dedicated to an analysis of the *Lophophytum mirabile* (also a completely heterotrophic plant from the same family as *R. phalloides*) did not observe contigs with plastid genes (Sanchez-Puerta et al., 2017). Additionally, in *R. lagascae*, which was reported to have no plastid genome (Molina et al., 2014), it may potentially be present but be unnoticed due to its high AT content. *Rafflesia lagascae* genome assembly was performed using circa 400 million Illumina reads, the same amount as we used for the assembly of *R. phalloides*, and thus if *R. lagascae* indeed possesses a plastid genome, it should be much more AT-rich than the plastid genome of *R. phalloides*.

The AT content is high in protein-coding genes (the average value weighted by length is 88.1%), as well as ncRNA-coding genes (the average value weighted by length is 77.5%) and non-coding regions (the average value weighted by length is 93.8%). In protein-coding genes, this led not only to a shift in codon frequencies towards AT-rich codons (Supplementary Table 2) but also to a shift in amino acid frequencies in proteins, with amino acids encoded by AT-rich codons being more used (Figure 3, Supplementary Table 3). For example, isoleucine, the amino acid with the most AT-rich codons, is used 2 times more often in the proteins encoded in the plastid genome of *R. phalloides* than in homologous proteins of phylogenetically close mixotrophic species. Similarly, glycine, whose codons are among the most GC-rich, is used 2 times more rarely.

**Figure 3.**
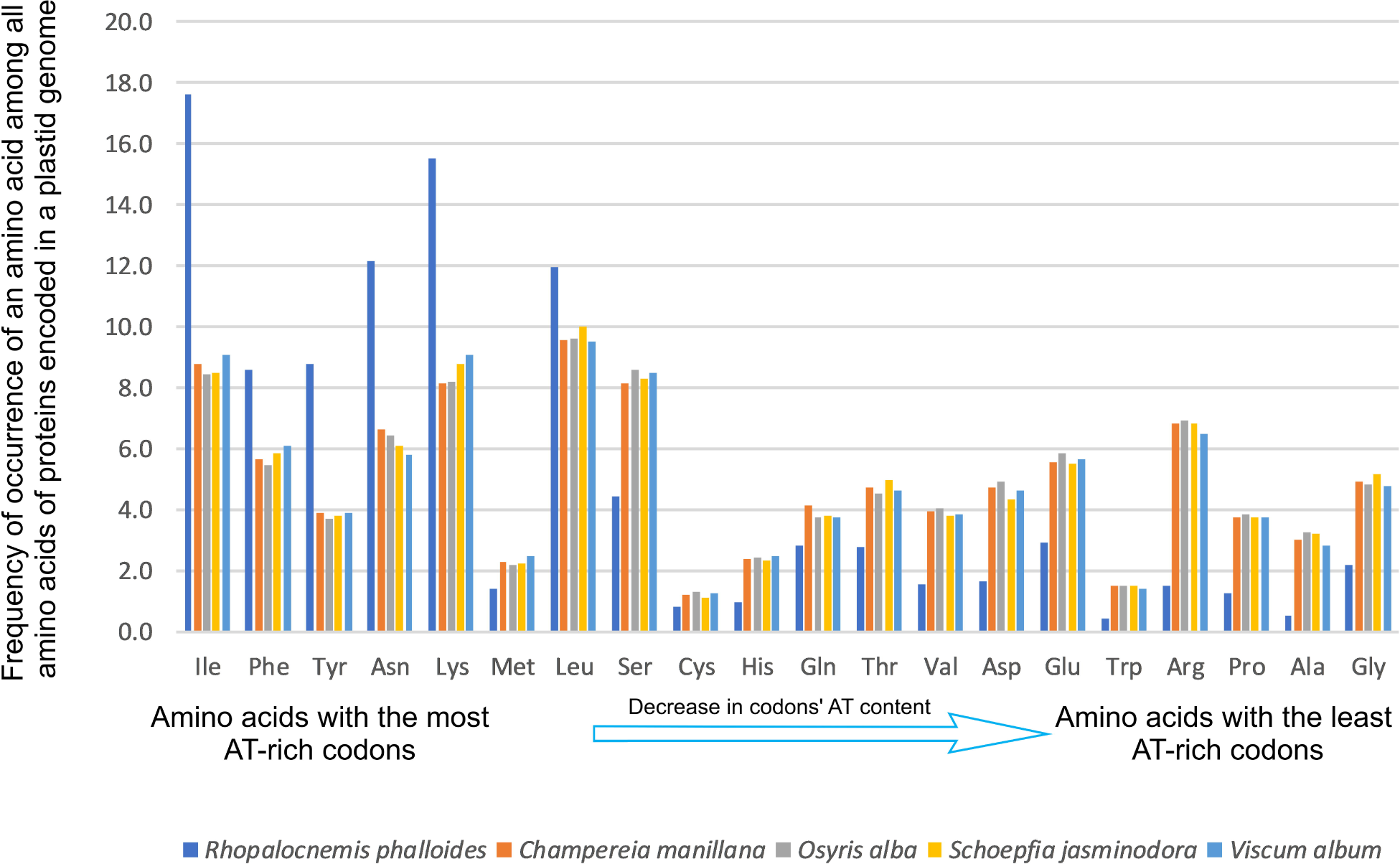
Amino acid usage in the plastid proteins of *Rhopalocnemis phalloides* is affected by the high AT content.

Interestingly, such high AT content has led to convergence of gene sequences of *R. phalloides* with sequences from phylogenetically distant AT-rich species. When aligning sequences of genes and proteins of *R. phalloides* to sequences from NCBI NT and NR databases, respectively, the best matches are often sequences from distantly related heterotrophic plants whose plastid genomes also have high AT content (Supplementary Table 4). There are also many matches to sequences from various protists and some matches to sequences of animals and bacteria. Not all the matches are to homologous sequences, with some resulting from accidental similarity to non-coding sequences.

We thoroughly investigated one of the prominent cases of convergence—the *rrn16* gene. BLASTN alignment of *rrn16* of *R. phalloides* to NCBI NT produces two best hits to other species of Balanophoraceae, namely *C. crassa* and *B. japonica* (for which only this plastid gene is sequenced and available in GenBank), while the next several dozen matches are to protists from the genera *Plasmodium*, *Nitzschia* and *Leucocytozoon*, belonging to SAR. Our initial hypothesis was a horizontal transfer from SAR to a common ancestor of the aforementioned Balanophoraceae. This is supported by the fact that a phylogenetic analysis of *rrn16* places the sequences of Balanophoraceae within SAR with bootstrap support value of 100 (Figure 4). A simple counterargument is that *Plasmodium*, *Nitzschia* and *Leucocytozoon*, though all belonging to SAR, are, in fact, quite distant phylogenetically from each other (with *Nitzschia* belonging to Stramenopiles, and *Plasmodium* and *Leucocytozoon* to Alveolata), and thus the fact that they appear in BLAST results together suggests some sort of bias. What is common for the species of the genera whose *rrn16* produces best matches to *rrn16* of *R. phalloides* is that they have extremely high AT content, close to that of *R. phalloides*. This led us to guess that the similarity originates not from the phylogenetic relatedness of *rrn16* of *R. phalloides* to *rrn16* of these species but from convergence because of their high AT content. To verify this, we rebuilt the phylogenetic tree, excluding from the multiple alignment all columns with A or T in *R. phalloides*, thus eliminating the possible convergence originating from the high AT content. In the resulting tree, *R. phalloides* is placed among plants (Figure 4C), which is correct, confirming that the placement in SAR was due to the AT content. Removal of columns with A and T in *B. japonica* or *C. crassa* leds to similar results: in the produced trees (Supplementary Figure 3) these species are situated either within plants (in the case of *B. japonica*) or between plants and SAR (in the case of *C. crassa*). An alternative explanation for the seeming phylogenetic closeness of *rrn16* of these three species of Balanophoraceae to *rrn16* of SAR can be the long branch attraction, but it is a characteristic problem of the maximum parsimony method, and it affects phylogenetic trees built with the maximum likelihood method to a lesser degree (Kück et al., 2012). Additionally, the similarity of *rrn16* orthologues can potentially be a result of misalignment, but the alignment is good, and the convergence is clearly observed in the alignment (Supplementary Figure 4).

**Figure 4.**
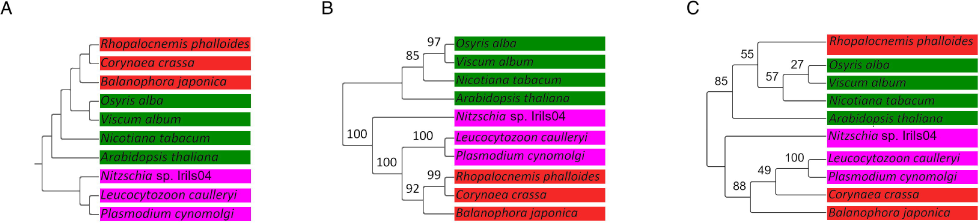
*rrn16* of *Rhopalocnemis phalloides* shows convergence with *rrn16* from SAR due to high AT content. (A) Phylogenetic tree of species. (B) Phylogenetic tree of *rrn16*. (C) Phylogenetic tree of *rrn16*, built by alignment columns where *Rhopalocnemis phalloides* has guanine or cytosine. Species with names in red rectangles are non-photosynthetic plants from Balanophoraceae, species with names in green rectangles are photosynthetic plants, and species with names in purple rectangles are from SAR. The numbers on the branches are bootstrap support values. The second and the third tree are unrooted.

Overall, our results suggest that phylogenetic analyses of heterotrophic plants (and, in general, of any species whose genomes have highly biased nucleotide composition) should be performed cautiously, as even bootstrap support values of 100 do not guarantee reliable phylogenetic reconstruction in such cases.

### Natural selection and substitution rate in the plastid genome of *R. phalloides*

Nucleotide substitution rate is known to be increased in plastid genomes of heterotrophic plants, ranging from a hardly detectable increase in plants that have lost their photosynthetic ability recently (Barrett et al., 2018) to a nearly 100-fold increase with respect to the closest photosynthetic species in the most reduced plastid genomes (Bellot and Renner, 2015). The reason for this increase is, to our knowledge, not yet known (and will be discussed in more details in the section “Why is the AT content so high?”).

To compare the substitution rate in *R. phalloides* with rates in its closest mixotrophic relatives, one should first determine the phylogenetic placement of *R. phalloides* relative to the species of comparison. The placement of the family Balanophoraceae has long been debated, with some scientists stating that it does not even belong to Santalales (Cronquist, 1981; Kuijt, 1968; Takhtadzhi□a□n, 2009). A recent work, which utilized sequences of 7 genes for phylogeny evaluation, suggests that Balanophoraceae indeed belong to Santalales (Su et al., 2015). More, the results of that work suggest polyphyly of Balanophoraceae, which consists of two clades: “Balanophoraceae A” and “Balanophoraceae B”. A common feature of Balanophoraceae A is that they have highly increased substitution rates, and a common feature of Balanophoraceae B is that their substitution rates are approximately the same as in autotrophic and mixotrophic Santalales. Although it analysed 11 species of Balanophoraceae, that study did not analyse *R. phalloides*. To estimate the phylogenetic relationships of *R. phalloides*, we added its data to the alignment of sequences from 186 species used in that article and rebuilt the tree. As one may have expected, *R. phalloides* is placed in Balanophoraceae A, with a bootstrap support value of 100 (Supplementary Figure 5). It is sister to a group of *C. crassa* and *Helosis cayennensis*.

To evaluate substitution rates, dN and dS in the plastid genome of *R. phalloides*, we used common protein-coding genes of this genome and plastid genomes of several other species of Santalales, available as of 2017. The genes *ycf1*, *ycf2* and *rps7* were excluded from the analysis because due to the high amount of accumulated mutations, their sequences in *R. phalloides* cannot be reliably aligned with homologous sequences of other species. The analysis by PAML shows that the number of nucleotide substitutions in the plastid genome of *R. phalloides* since the divergence from common ancestor with the mixotrophic Santalales of comparison is, on average, 264 times higher than in the plastid genomes of those mixotrophic Santalales (Figure 5). This number should be treated with caution, as:

**Figure 5.**
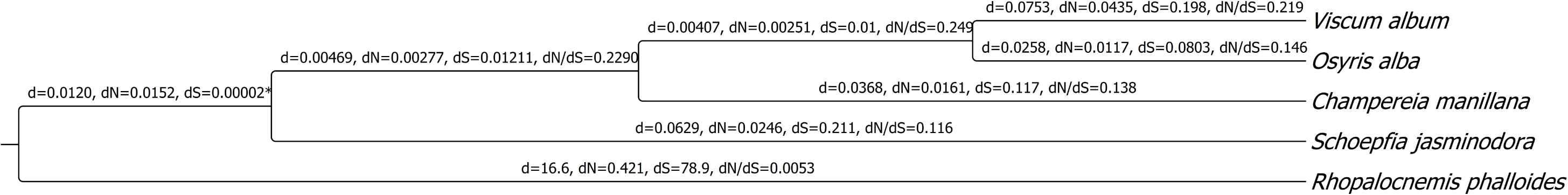
Evolutionary parameters of the phylogeny of Santalales. *Arabidopsis thaliana*, used as the outgroup, is not shown. “d” is the number of substitutions per nucleotide. The total length of the alignment, used for the analysis, was 4095 bp after removal of poorly aligned regions by Gblocks. * dN/dS on this branch cannot be calculated due to a very small dS value.

1. The model of nucleotide substitutions used in PAML utilizes the equilibrium codon frequencies, equal for all branches. This is definitely not the case in the studied Santalales, as the codon frequencies in the plastid genome of *R. phalloides* highly differs from those in plastid genomes of mixotrophic Santalales (Supplementary Table 2). We are aware of a single tool for phylogenetic analyses that is able to take into account different codon frequencies in different sequences. This is the program collection BppSuite. However, the analysis of these data by BppSuite provides the value of approximately 25 000 instead of 264, which is likely an algorithmic mistake.
2. Non-synonymous substitutions quickly reach saturation, and thus the number of non-synonymous substitutions is underestimated for long branches (dos Reis and Yang, 2013). The same is true for synonymous substitutions (Vanneste et al., 2013).
3. We removed columns in the alignment with many differences between species using the program Gblocks, since such columns may result from misalignment. As regions of genes with positive or weak negative selection accumulate mutations faster, such regions can also be potentially removed by Gblocks, leading to underestimation of substitution rates.
4. We failed to produce reliable alignments for the genes ycf1, ycf2 and rps7, consequently the substitution rate in these genes may be higher than in others. Therefore, the exclusion of these genes from the analysis may lead to underestimation of the true substitution rate.

While dN/dS values in the plastid genomes of the studied mixotrophic Santalales provided by PAML are typical (approximately 0.1-0.2), the value in *R. phalloides* is much lower, approximately 0.005. The underestimation of dN/dS on long branches is a known effect, which results from saturation in non-synonymous positions (dos Reis and Yang, 2013). Therefore, the actual dN/dS of the protein-coding genes in the plastid genome of *R. phalloides* is hard to estimate. The selection acting on the genes of *R. phalloides* is definitely non-neutral, as open reading frames of all the genes are intact. If we denote by P(X) the probability that there is a specific codon in a specific position, and by α the AT content of a gene, then the probability that a random codon is stop P(Stop)=P(TAA)+P(TGA)+P(TAG)=(α/2)×(α/2)×(α/2)+(α/2)×((1-α)/2)×(α/2)+(α/2)×(α/2)×((1- α)/2)=α^2^/4-α^3^/8.

As the weighted (by length) average AT content in protein-coding genes of *R. phalloides* is 88%, the probability of a random codon being stop, as follows from this equation, is approximately 11%. This means that since stop codons are AT-rich, in a random sequence with such a high AT content as in *R. phalloides*, every 9th codon will be stop. Therefore, a strong negative selection shall act on the genes to keep open reading frames unbroken.

### Other genomes of *R. phalloides*

Sequencing of circa 400 million paired-end reads could have been enough to assemble the mitochondrial and the nuclear genomes of *R. phalloides*. Alignment by BLASTN and TBLASTN of, respectively, ncRNAs and proteins encoded in mitochondrial genomes of reference species to the contigs of *R. phalloides* revealed several dozen matching contigs with coverages of approximately 5000× and length of approximately 1000-5000 bp. They are probably short mitochondrial chromosomes, similar to the ones observed in the plant *L. mirabile* (Sanchez-Puerta et al., 2017), also from Balanophoraceae, whose mitochondrial genome putatively consists of 54 small circular chromosomes. We do not plan to investigate the mitochondrial genome of *R. phalloides* in detail and are ready to provide the mitochondrial contigs by request.

Known sizes of nuclear genomes of plants from Santalales vary from approximately 200 Mbp in *Santalum album* (Mahesh et al., 2018) to approximately 100 Gbp in *Viscum album* (Zonneveld, 2010). If the nuclear genome size in *R. phalloides* is, for example, 500 Mbp, 400 million 150-bp-long reads will produce a coverage of approximately 400×150/500=120×, which is enough for a draft assembly. To estimate the nuclear genome size, we built a k-mer frequency histogram (Supplementary figure 6). The peak of the distribution, corresponding to the k-mer coverage of the nuclear genome, is hard to determine, but it is below the k-mer coverage value of 2×. As the k-mer size (21) is much lower than the read size (150), the read coverage is approximately equal to the k-mer coverage. Therefore, the nuclear genome size may be estimated as being at least 400×150/2=30 000 Mbp. Potentially, the genome size may be overestimated if there is a lot of contamination (for example by DNA of endophytic bacteria and fungi), but a taxonomic analysis of reads suggests that contamination in unlikely to be big (Supplementary Table 5). Assembly of a 30 000-Mbp-long genome is impossible using only the reads produced in the current study. Instead of the complete nuclear genome assembly, we plan to study it by means of transcriptome assembly, which is a subject of our next work.

### Why is the AT content so high?

The increase in the AT content in the plastid genomes of heterotrophic plants, as well as the increase in their substitution rates, are known and many times discussed phenomena (Bromham et al., 2013; Wicke et al., 2016; Hadariová et al., 2018; Wicke and Naumann, 2018). However, their origin is still unknown. The simplest hypothesis for the increase in the substitution rate may be the relaxation of selection acting on genes. However, plastid genes of heterotrophic plants usually show no signs of relaxed selection, except for photosynthesis-related genes during pseudogenization. Interestingly, the high AT content and substitution rate are also observed in plastids of non-photosynthetic protists (such as *Plasmodium*) (Oborník et al., 2009), which lost the genes required for photosynthesis after the transition to heterotrophic lifestyle. Additionally, both of these phenomena are found in genomes of endosymbiotic bacteria (McCutcheon and Moran, 2011), which may be dozens of times shorter than genomes of their free-living relatives due to the loss of genes required, for example, for biosynthesis of substances which are now provided to the symbiont by its host. Therefore, these two phenomena are probably not only unrestricted to plants but are even not related to the loss of photosynthesis.

A phenomenon that can simultaneously result both in increase of AT content and in increase of substitution rate is the reduction in genome recombination intensity. A plastid genome is capable of recombining both within itself (the recombination of two copies of the inverted repeat) (Li et al., 2016; Zhu et al., 2016) and between two copies of a genome (Maréchal and Brisson, 2010). The recombination is an important step in reparation, both in plastids (Zampini et al., 2017) and in bacteria (Cox, 1998), so the reduction in recombination will increase the substitution rate. Also, gene conversion in plastid (Wu and Chaw, 2015; Zhitao Niu et al., 2017) as well as in bacterial (Lassalle et al., 2015) genomes is GC-biased, although earlier gene conversion in plastid genomes was supposed to be AT-biased (Khakhlova and Bock, 2006). This means that if, during a recombination, there is a mismatch between an adenine or a thymine on one strand versus a guanine or a cytosine on the other, it is more likely that the guanine or the cytosine will be kept, while the adenine or the thymine will be removed and replaced by a cytosine or a guanine, which is complementary to the base on the other strand. Therefore, recombination aids in increasing GC content in plastid and bacterial genomes, and a decrease in recombination will make a genome more AT-rich. The link between the low recombination rate and the high AT content has already been proposed for endosymbiotic bacteria with small genomes (Lassalle et al., 2015).

Recently, it was shown that in transcriptomes of the heterotrophic plants *Epipogium aphyllum*, *E. roseum* and *Hypopitys monotropa*, the transcript of the protein RECA1, required for recombination of the plastid genomes, is absent (Schelkunov et al., 2018). This may support the above hypothesis. However, the direct reason for the loss of RECA1 is not known. A potential explanation for the loss may be that during the transition from a mixotrophic to a heterotrophic lifestyle, plastid enzymes related to photosynthesis accumulate mutations, and since a mutated enzyme may be harmful for the organism, it is evolutionarily adaptive to accumulate the mutations very fast to quickly achieve complete disruption of a gene, instead of having a semi-crumbled gene encoding a harmful protein. This effect, consisting in elimination of pseudogenes with rates faster than neutral, was already shown to take place in bacteria (Kuo and Ochman, 2010). Therefore, in the period directly following the loss of photosynthesis, it may be beneficial for the plant to disturb the plastid recombination and thus disturb the repair. In fact, this process may start even before the loss of photosynthesis, since in plastid genomes of mixotrophic plants *ndh* genes often undergo pseudogenization (Wicke et al., 2011), and their quick removal may require an increased mutation accumulation rate. Such an increase in the mutation accumulation rate may require pseudogenization of genes of DNA replication, recombination and repair (DNA-RRR), such as *RECA1*, and once they are pseudogenized, it will be hard for a plant to return to the normal repair intensity in the plastid genome, thus making the transition to high mutation accumulation rates irreversible.

It is known that the mutation accumulation rate in heterotrophic plants (Bromham et al., 2013), including Balanophoraceae (Su and Hu, 2012; Su et al., 2015), is also increased in nuclear and mitochondrial genomes, although to a lesser extent than in plastid genomes. These phenomena are also still unexplained. The nuclear genome contains more than a hundred (Schelkunov et al., 2018) genes that encode proteins, working in multisubunit complexes with proteins, encoded in the plastid genome. These are the genes encoding proteins of the electron-transfer chain, the plastid-encoded RNA polymerase (PEP), the plastid ribosome and others. When a species loses its photosynthetic ability, the nuclear-encoded genes of the electron-transfer chain are no longer under selective pressure and start to accumulate mutations. Therefore, their proteins may become harmful and may require quick elimination. Thus, the increase in the nuclear mutation accumulation rate, which may speed up the accumulation of disruptive mutations in these genes, may also by selectively beneficial. The increase in the mutation accumulation rate in the mitochondrial genome may be potentially explained by the fact that many DNA-RRR proteins are common for the plastid and the mitochondrial genomes (Shedge et al., 2007; Carrie and Small, 2013). Therefore, if it is selectively beneficial to increase the mutation accumulation rate in the plastid genome, the mitochondrial genome may also be affected.

This hypothesis of accelerated junk removal may be tested by studying plastid and nuclear genomes of many related heterotrophic species and checking whether the crumbling genes accumulate mutations with rates faster than neutral shortly after the loss of photosynthesis and whether some of the DNA-RRR genes deteriorate at the same time.

## Conclusions

The plastid genome of *R. phalloides* profoundly differs from plastid genomes of typical plants, including the massive gene loss, the increased substitution rate and the high AT content. By decreasing sequencing coverage, such high AT content may “hide” plastid genomes of some heterotrophic plants, making these genomes harder to find by means of high-throughput sequencing. Alterations in the nuclear genome, accompanying these changes in the plastid genome, are an interesting issue. Our next work will be dedicated to the study of the nuclear genome of *R. phalloides* by means of transcriptome sequencing.

## Acknowledgements

The work was funded by a Russian Foundation for Basic Research grant (No. 16-34-01003). The work of MSN was carried out in accordance with a Government order for the Lomonosov Moscow State University (project No. AAAA-A16-116021660105-3). We are grateful to Alina Alexandrova for field assistance.

## Data accessibility

The plastid genome is deposited in the GenBank database under accession MK036331. The DNA-seq and RNA-seq reads are deposited in the Sequence Read Archive (SRA) under accessions SRR7995544 and SRR7995545, respectively.

## Figures and tables

**Supplementary Figure 1. Dependence of the sequencing coverage on the AT content in the plastid genome of *Rhopalocnemis phalloides***. The sequencing coverage is provided on a logarithmic scale. The window size for the calculation was 241 bp (the average insert size). The step for window sampling was 50 bp.

**Supplementary Figure 2. Dependence of the insert size on the AT content in the plastid genome of *Rhopalocnemis phalloides***. The window size for the calculation was 241 bp (the average insert size over the whole genome). The step for window sampling was 50 bp.

**Supplementary Figure 3. Influence of AT-rich positions on the phylogeny of *rrn16***. (A) Phylogenetic tree of *rrn16*, built by alignment columns where *Balanophora japonica* has guanine or cytosine. (B) Phylogenetic tree of *rrn16*, built by alignment columns where *Corynaea crassa* has guanine or cytosine. Species with names in red rectangles are non-photosynthetic plants from Balanophoraceae, species with names in green rectangles are photosynthetic plants, and species with names in purple rectangles are from SAR. The numbers on the branches are bootstrap support values. The trees are unrooted.

**Supplementary Figure 4. Alignment of *rrn16* demonstrates convergence between Balanophoraceae and SAR in AT-rich positions.** Columns removed by Gblocks are shaded. These columns mostly correspond to regions of SAR that were not sequenced, as Gblocks removes all columns that have gaps.

**Supplementary Figure 5. Phylogenetic tree of Santalales.** Numbers over branches denote bootstrap support values. *Antirrhinum majus*, *Cornus florida*, *Camellia japonica*, *Spinacia oleracea*, *Myrtus communis* and *Arabidopsis thaliana* are used as the outgroup. The scale bar represents the number of nucleotide substitutions per position.

**Supplementary Table 1. Genome lengths and AT content among the plastid genomes of Embryophyta.** Names of completely heterotrophic species are coloured orange, and names of mixotrophic or completely autotrophic are coloured green.

**Supplementary Table 2. Codon usage in the plastid genomes of Santalales.** Data corresponding to *Rhopalocnemis phalloides* are coloured orange, and data corresponding to mixotrophic Santalales are coloured green. Numbers denote percentages of codons used to encode a specific amino acid, compared to all codons of this amino acid.

**Supplementary Table 3. Amino acids usage in the proteins encoded in the plastid genomes of Santalales.** Data corresponding to *Rhopalocnemis phalloides* are coloured orange, and data corresponding to mixotrophic Santalales are coloured green. Numbers are percentages, representing an amount of a specific amino acid compared to the total number of amino acids.

**Supplementary Table 4. BLAST matches between genes and proteins of *Rhopalocnemis phalloides* and genes and proteins from NCBI NT and NCBI NR databases.**

**Supplementary Table 5. Taxonomic distribution of best BLAST matches for 1000 random read pairs.** The read pairs with no matches likely correspond to huge non-coding regions of Rhopalocnemis phalloides’ nuclear genome.

